# Forest maturity and functional nestedness shape harvestman trait diversity in the Atlantic Forest

**DOI:** 10.64898/2026.06.10.731358

**Authors:** Rick Chalela Curdoglo, Lívia Santos Lourenço, Beatriz Kali Silva, Stefan Ribeiro, Cibele Bragagnolo

## Abstract

Tropical forest succession can reorganize biodiversity not only by changing species richness, but also by filtering the functional traits that represent ecological strategies. Harvestmen are highly sensitive to microclimatic and structural changes in Neotropical forests, yet their functional diversity remains poorly explored. We investigated taxonomic and functional diversity of harvestmen in a local scale, an Atlantic Forest remnant in southeastern Brazil containing forest patches at different successional stages. Standardized nocturnal active searches and leaf-litter sampling yielded 384 individuals belonging to 14 morphospecies. Functional diversity was quantified from four morphological traits using Hill numbers (q = 0, 1 and 2), morphofunctional ordination, beta-diversity partitioning and environmental models based on forest-structure variables and PCA-derived gradients. Functional diversity was highest when rare species were weighted equally and declined strongly from q = 0 to q = 2, indicating that uncommon species carried much of the regional morphofunctional variation. Functional alpha diversity was positively associated with taxonomic alpha diversity, and Mantel tests showed that taxonomic and functional dissimilarities among sampling points were significantly correlated. However, formal beta-diversity partitioning refined this interpretation: functional beta diversity was dominated by nestedness-resultant dissimilarity rather than turnover, suggesting that functionally poorer assemblages represented contracted subsets of the regional trait space. Morphofunctional analyses identified compact, robust and long-legged species groups, and environmental models showed that lower vegetation structure, litter depth and forest-maturity gradients significantly influenced functional diversity. These findings indicate that mature, structurally complex Atlantic Forest patches help maintain the full spectrum of harvestman morphofunctional strategies and highlight harvestmen as promising models for trait-based conservation ecology.

**Graphical Abstract:** 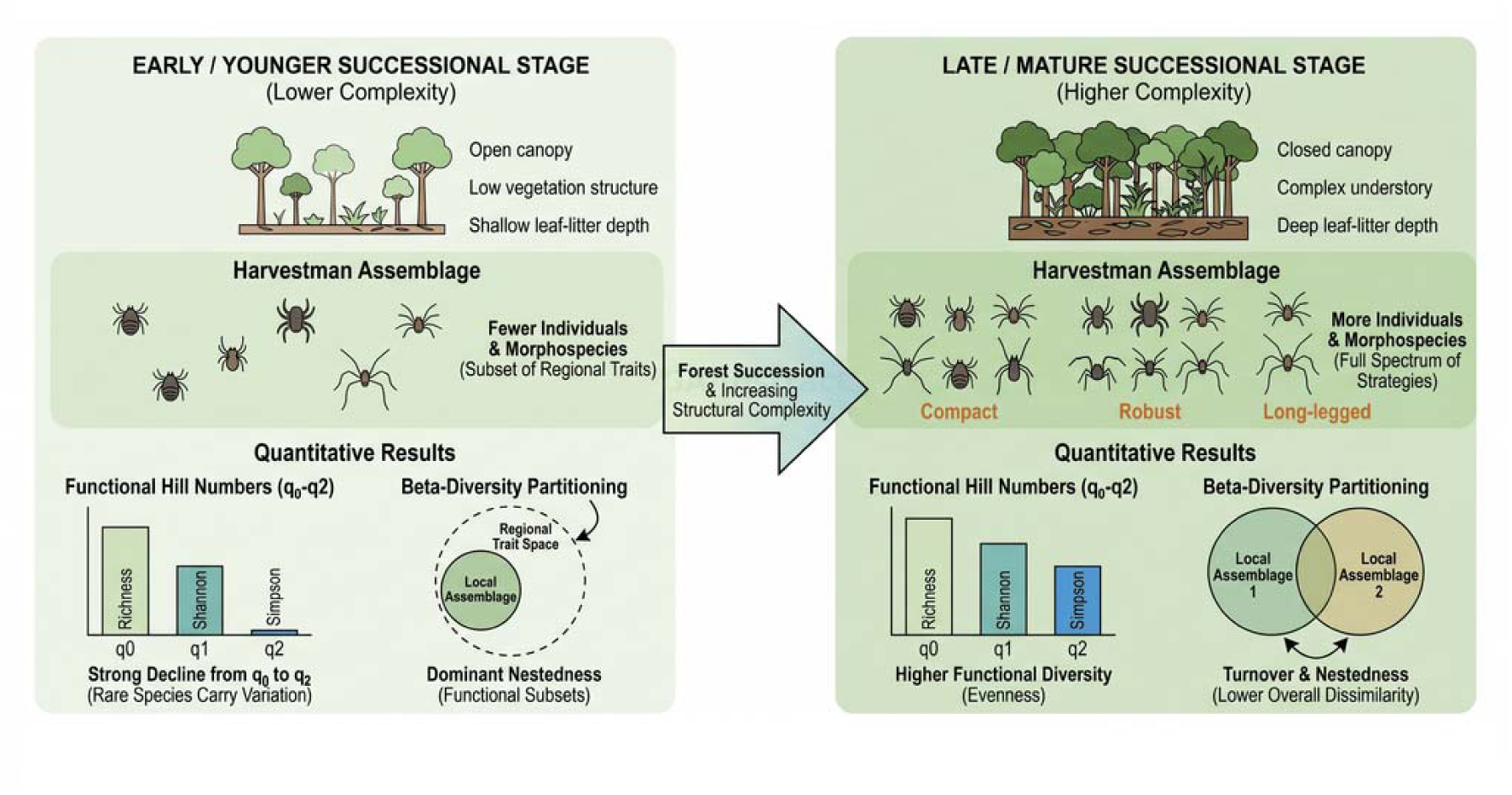

## 1. Introduction

Biodiversity change is commonly described through the number of species occurring within local communities, across landscapes and among habitats, in different scales [6]. This taxonomic perspective remains essential for ecology and conservation, but it captures only part of the biological variation that determines how communities respond to environmental change [14,10]. Alpha diversity describes the diversity found locally, gamma diversity represents the diversity accumulated across a broader region, and beta diversity expresses changes in community composition along spatial or environmental gradients, including turnover and nestedness processes [19,15].

During recent decades, community ecology has increasingly incorporated additional dimensions of biodiversity, especially functional and phylogenetic diversity [46,29]. Functional diversity is particularly useful because it links species occurrence to measurable traits that can mediate performance, microhabitat use, dispersal ability, trophic interactions and sensitivity to environmental filters [48,44]. Communities with high functional complementarity may contain species that use different resources or microhabitats, whereas low functional redundancy may indicate that the loss of even rare species can remove unique ecological strategies [29,22].

Recent studies have emphasized that functional diversity should be evaluated not only through local alpha components, but also through functional dissimilarity among communities [39]. By partitioning beta diversity into turnover and nestedness-resultant components, it becomes possible to distinguish whether differences among sites are mainly driven by the replacement of species or functional strategies, or by the loss or gain of subsets of the regional functional space. For example, Stefanidis [39] showed that comparing taxonomic and functional beta diversity can reveal whether environmental gradients structure communities through species replacement, functional replacement or functional nestedness. This framework is especially relevant for habitat-sensitive organisms, because structurally distinct habitats may support different combinations of functional traits even when taxonomic richness alone shows limited variation.

This functional perspective is especially relevant in tropical forests, where environmental heterogeneity, vertical vegetation structure and microclimatic stability can generate fine-scale niche differentiation [34]. In fragmented or regenerating Atlantic Forest landscapes, structural simplification may therefore reduce not only taxonomic diversity but also the range of ecological strategies supported by the habitat [3,24,4].

Among terrestrial arthropods, functional diversity has been more frequently explored in insects and spiders than in other arachnid groups [38,33]. Harvestmen remain neglected from a functional-trait perspective despite their ecological importance in Neotropical forests. The Atlantic Forest is one of the major centers of harvestman diversity [30,41] and besides these animals are sensitive to habitat disturbance and fragmentation [21,30,31], little is known about how their morphological traits vary across local communities and environmental gradients [32].

Here, we investigated taxonomic and functional diversity patterns of harvestmen at a local scale in a continuous Atlantic Forest remnant in southeastern Brazil, where forest patches differ in regeneration age and structural conditions. We tested three main hypotheses: (1) functional alpha diversity increases with taxonomic alpha diversity, indicating low functional redundancy among local assemblages; (2) taxonomic dissimilarity among sampling points is positively associated with functional dissimilarity, suggesting that changes in species composition are accompanied by changes in morphofunctional composition; and (3) mature and structurally complex forest patches support higher functional diversity and a more complete representation of the regional functional trait space than younger or structurally simplified patches. We further partitioned taxonomic and functional beta diversity into turnover and nestedness-resultant components to determine whether among-site differences were mainly driven by replacement of species and functional strategies or by nested losses of taxonomic and functional diversity. By focusing on an understudied arachnid group in a highly threatened tropical forest, we aim to strengthen the use of harvestmen as models for functional biodiversity and conservation ecology.

## 2. Methods

### 2.1. Study area

The study was conducted at Parque Ecológico Imigrantes (PEI), a private protected area located near Serra do Mar in São Bernardo do Campo, São Paulo state, southeastern Brazil (23°50’47.81"S, 46°34’42.44"W; Fig. 1). The park comprises approximately 484,000 m² (48.4 ha) of dense ombrophilous Atlantic Forest at approximately 772 m a.s.l. Mean annual temperature ranges from approximately 14.5 °C in winter to 21 °C in summer, and mean annual rainfall is approximately 1,524 mm [8]. Parts of PEI were historically affected by deforestation, resulting in a mosaic of forest patches at different regeneration stages. Based on georeferenced aerial images from 1962, 1973, 1980 and 2020, the park was classified into forest-age categories that allowed harvestman communities to be evaluated along a local gradient of forest structural development.

**Figure 1.**
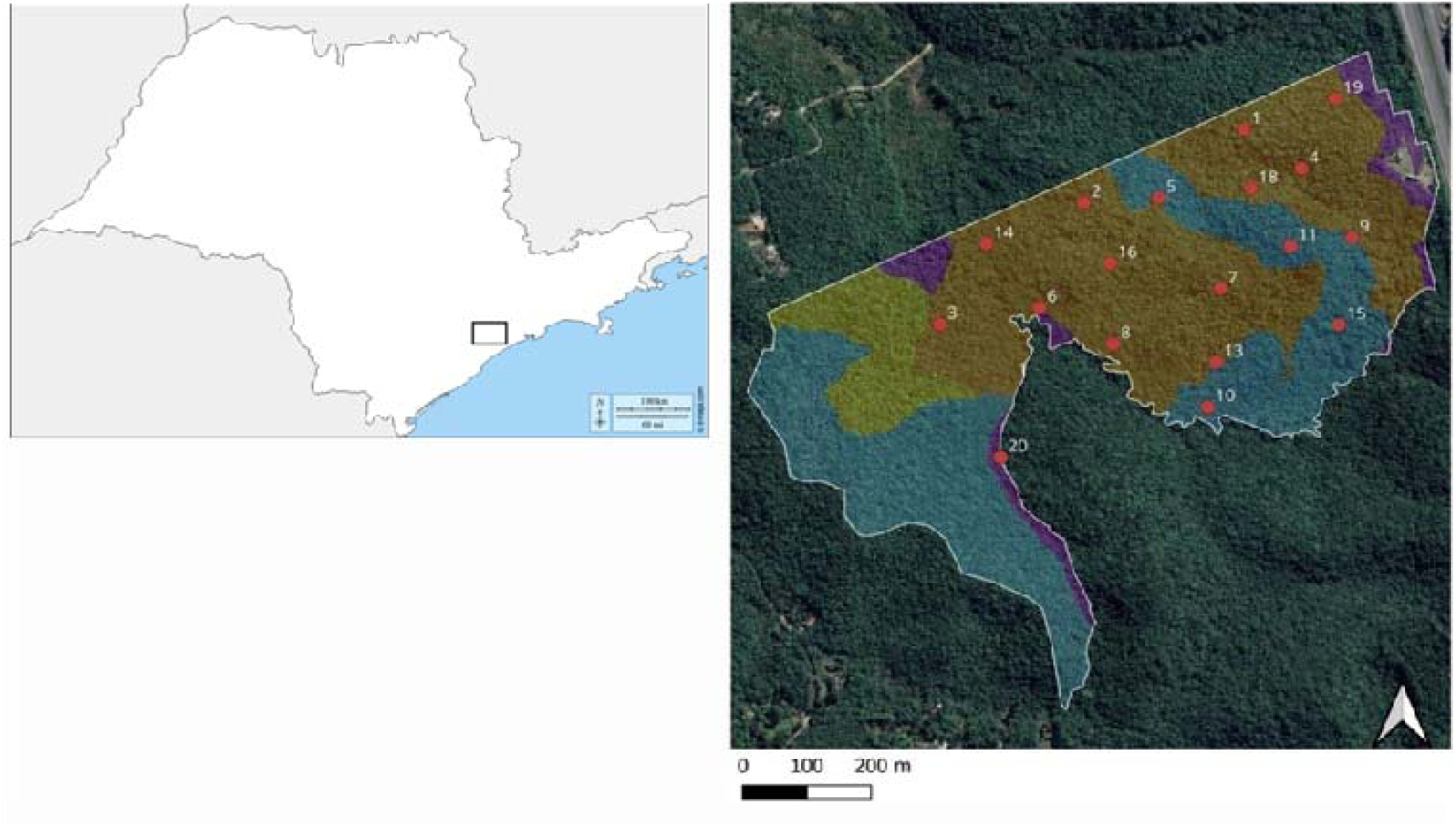
Representative map of the Parque Ecologico dos Imigrantes (PEI) area. Red circles: 18 sampling points; dashed White line: PEI property boundaries; colored polygons represent forest areas of varying ages, with blue representing forests over 60 years old, yellow representing forests approximately 47 years old, orange representing forests approximately 40 years old, and purple representing forests less than 40 years old. Maps data: Google, ⓒ2025 Airbus.

### 2.2. Sampling design and harvestman collection

Sampling points were initially selected at random, with a minimum distance of 100 m among points. Sampling points were positioned at least 50 m from forest edges to reduce edge effects and at least 30 m apart to avoid sampling overlap. Due to logistical restrictions related to PEI supervision, some initially selected points could not be sampled using all methods. Therefore, the final sampling effort comprised 54 nocturnal active-search samples and 36 leaf-litter samples across 18 sampling points (Fig. 1).

Harvestmen were sampled using two complementary methods: standardized nocturnal active search and leaf-litter sampling. These methods were used to sample species associated with different microhabitats, including understory vegetation, tree trunks, stones and the leaf-litter layer.

Standardized nocturnal active searches were performed using headlamps, inspecting substrates from ground level up to accessible vegetation height, including trunks, fallen branches and other microhabitats. Observed arachnids were captured, placed in vials and immediately preserved in 70% ethanol. Each sampling unit consisted of one collector searching for 1 h along a 30 m transect, inspecting substrates within approximately 5 m on each side of the transect. At each of the 18 sampling points, three collectors sampled simultaneously, resulting in three standardized sampling units per point and 54 active-search samples in total.

Leaf-litter sampling was performed using a 0.5 × 0.5 m sampling frame. At each selected point, leaf litter within the frame was removed with a shovel and placed in plastic bags for subsequent sorting and inspection. Two leaf-litter samples were collected at each of the 18 sampling points, during the day time, one in February and one in August 2022, totaling 36 leaf-litter samples.

Field activities and the collection of zoological material were conducted in accordance with Brazilian environmental legislation. The corresponding author holds a permanent license for the collection of zoological material issued through the Brazilian Biodiversity Authorization and Information System (SISBIO), which authorized the collection, transport and preservation of the specimens used in this study.

### 2.3. Sorting and taxonomic identification

Collected material was transported to the Arachnology Laboratory of Universidade Federal de São Paulo. Specimens were sorted and separated into spiders and harvestmen. Harvestmen were morphotyped and, when possible, identified to the lowest feasible taxonomic level using taxonomic references for Opiliones [30] and a Leica M205C stereomicroscope. The biological material analyzed during this study is deposited in the Museu de Zoologia da Universidade de São Paulo (MZUSP).

### 2.4. Functional traits

Functional characterization was based on four morphological traits chosen to represent body size, body robustness and appendage length: (1) body length, measured from the base of the pedipalps to the posterior end of the body; (2) maximum body width at the level of coxa IV; (3) femur IV length; and (4) leg II length (Fig. 2). These traits were selected because they may be associated with locomotion, substrate use, exposure to desiccation and vertical microhabitat occupation.

**Figure 2.**
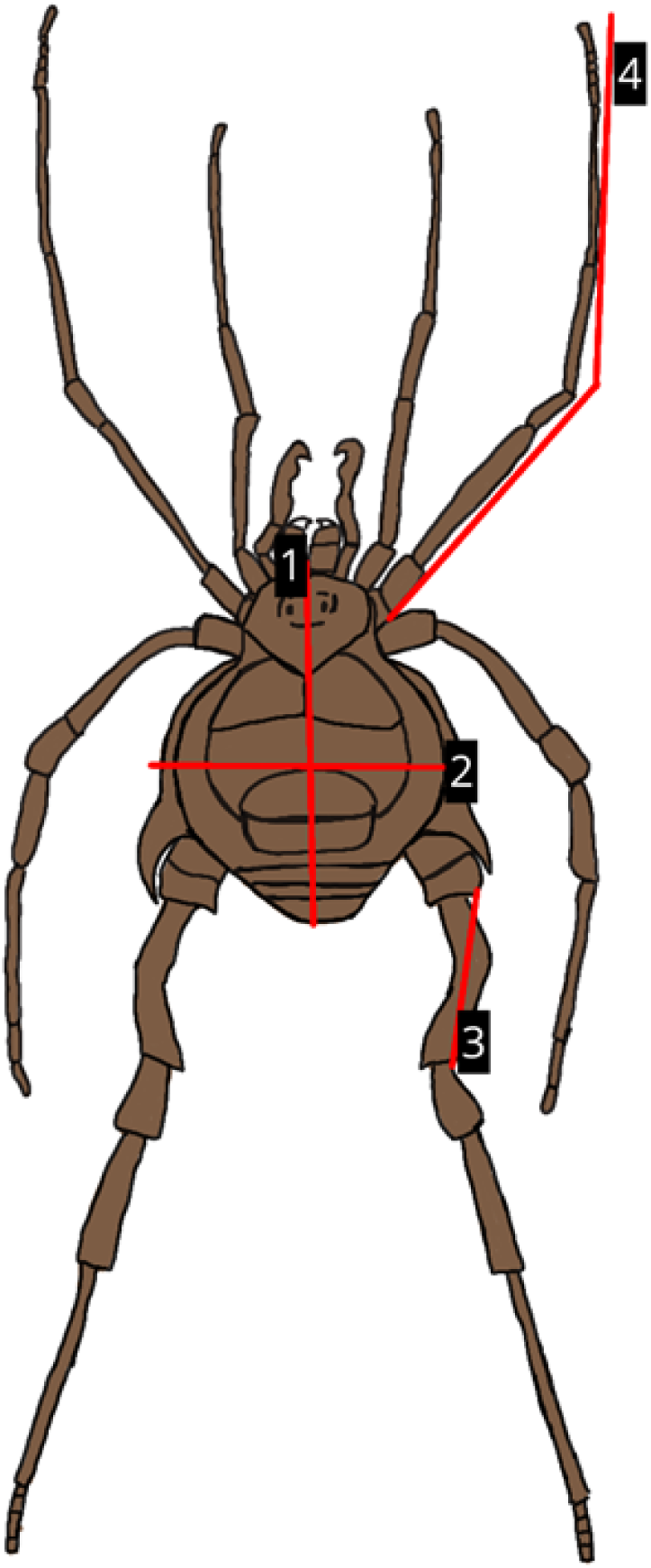
Morphological measurements used to characterize harvestman functional traits: body length, body width, femur IV length and leg II length.

Whenever possible, 20 individuals per species were measured, including 10 females and 10 males (see supplementary Table A2). For rare species with fewer available individuals, all collected individuals were measured. Measurements were taken using a digital caliper and a stereoscopic microscope.

### 2.5. Environmental variables

Environmental variables were selected to describe abiotic conditions and forest structure relevant to harvestman habitat use. For vegetation sampling, a 6 m diameter plot (approximately 28 m²) was delineated at the center of each sampling point, following the definitions of Higgins [13]. Within each plot, we measured variables that reflect habitat structure and quality:

- Diameter at breast height (DBH): trunk diameter at 1.4 m above ground for all trees with DBH ≥ 10 cm, measured by a single observer to standardize procedures.
- Canopy height: visually estimated with a 6 m extendable aluminum pole marked at 1 m intervals, using a random location within each plot.
- Vegetation stratification: assessed with the same pole following a vertical-structure protocol based on Pardini [27]. Four random measurements were taken per point,and foliage presence was recorded within a 1 m vertical column in five height bands: 0–1 m, 1–5 m, 5–10 m, 10–15 m and >15 m.
- Canopy cover: estimated from hemispherical photographs taken with a fisheye lens attached to a mobile phone. Photographs were taken in the morning on the same day in May 2022, and percentage foliage cover was quantified with ImageJ v.1.8 [37].
- Tree density: calculated as the number of trees with DBH ≥ 10 cm divided by the plot area.
- Forest age: classified from georeferenced aerial images from 1962, 1973, 1980 and 2020. Forest age was reclassified into two categories, younger secondary forest (< 40 years) and older/mature forest (> 40 years) (Fig. 1) for statistical analyses.
- Temperature: measured with a digital thermohygrometer at two random locations within each plot on different days, one in January and another in July. Analyses used the mean of the two measurements per point.
- Litter depth: measured with a 30 cm bamboo rod with a movable metal indicator; depth was measured with calipers in millimeters.

### 2.6. Statistical analyses

To reduce redundancy among environmental variables, pairwise Pearson correlations were calculated among quantitative forest-structure variables, and point-biserial correlations were used between forest age and quantitative variables. Correlated variables were summarized using principal component analysis (PCA), producing orthogonal axes for subsequent modeling (Figs. 3, 4).

**Figure 3.**
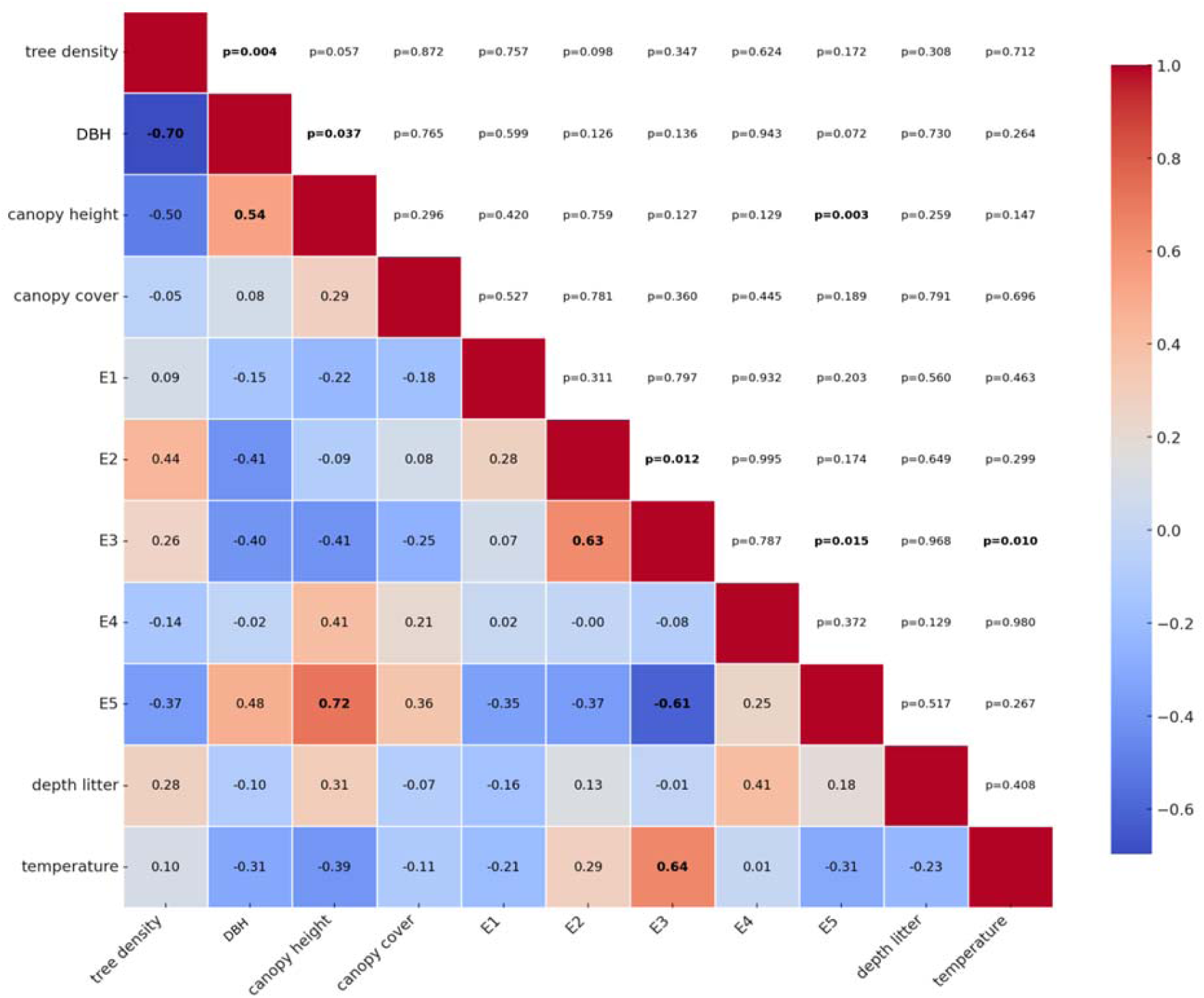
Correlation analysis (Pearson) between all pairs of vegetation structure variables. Lower quadrant indicates correlation strength, upper quadrant statistical significance. Statistically significant values are in bold (*p* < 0.05). E1–E5: vegetation stratification (0–1 m: E1, 1–5 m: E2, 5–10 m: E3, 10–15 m: E4, *>*15 m: E5). DBH: diameter at breast height.

Because sampling points were spatially distributed within the same continuous forest remnant, we evaluated whether environmental predictors and community response variables showed spatial structure. Spatial autocorrelation was assessed using Moran’s I based on the geographic coordinates of sampling points. This procedure allowed us to verify whether functional diversity patterns were primarily associated with forest-structure gradients rather than spatial proximity among sampling points.

Taxonomic alpha diversity was calculated as the observed number of harvestmen morphospecies recorded at each sampling point. Therefore, taxonomic alpha diversity corresponded to local morphospecies richness, equivalent to the taxonomic Hill number of order q = 0. This metric was used to test whether sampling points with higher species richness also supported higher functional diversity.

Functional diversity was calculated from two matrices: a species-by-site abundance matrix and a species-by-trait matrix. The abundance matrix contained the number of individuals of each harvestman morphospecies recorded at each sampling point. The trait matrix contained mean values per morphospecies for four continuous morphological traits: body length, maximum body width, femur IV length and leg II length. For morphospecies represented by multiple individuals, trait values were averaged across 20 measured individuals (10 males and 10 females); for rare morphospecies, all available individuals were measured, and their mean trait values were used.

Functional alpha, beta and gamma diversity were quantified using the Hill-number framework for functional diversity following Chiu [7]. Hill orders q = 0, q = 1 and q = 2 were used to represent different levels of abundance weighting. At q = 0, all species were weighted equally, emphasizing rare morphospecies. At q = 1, species were weighed according to their observed abundances, representing the functional diversity of typical species in the assemblage. At q = 2, abundant species received greater weight, emphasizing dominant morphospecies.

Functional alpha diversity represented the mean functional diversity within sampling points, whereas functional gamma diversity represented the total functional diversity of the assemblage across the study area. For functional diversity analyses, a pairwise functional distance matrix among morphospecies was calculated using Euclidean distances based on the standardized trait matrix, because all traits were continuous morphological measurements.

Functional beta diversity within the Hill-number framework was calculated multiplicatively as gamma diversity divided by alpha diversity. Thus, functional beta diversity expresses functional differentiation among sampling points for each Hill order q = 0, q = 1 and q = 2.

The relationship between taxonomic alpha diversity and functional alpha diversity was assessed using Spearman correlation. For this analysis, taxonomic alpha diversity was represented by observed morphospecies richness per sampling point, and functional alpha diversity was represented by functional Hill diversity at q = 0, because this order gives equal weight to all morphospecies and emphasizes the contribution of rare functional types.

Taxonomic and functional beta diversity were also evaluated using a dissimilarity-based framework to partition total beta diversity into turnover and nestedness-resultant components. This analysis was conceptually distinct from the multiplicative Hill-number beta diversity described above. For taxonomic beta-diversity partitioning, the species-by-site abundance matrix was converted to presence–absence data. Taxonomic dissimilarity was decomposed into total Sørensen dissimilarity (βSOR), turnover (βSIM) and nestedness-resultant dissimilarity (βSNE), following Baselga [1].

Functional beta-diversity partitioning was performed following Villéger [42], which decomposes functional dissimilarity into functional turnover and functional nestedness-resultant components.

We tested the relationship between taxonomic and functional dissimilarity using Mantel permutation tests. Mantel tests were performed between total taxonomic and functional dissimilarity matrices, and separately between their turnover and nestedness-resultant components, using 9,999 permutations. As a sensitivity analysis, we also compared abundance-based Bray–Curtis taxonomic dissimilarity with Euclidean distances among abundance-weighted community mean trait values.

To characterize morphofunctional structure, species mean trait values were standardized and summarized using PCA. A hierarchical clustering analysis using Ward’s method and Euclidean distances was used to define morphofunctional groups [45]. Differences among groups in multivariate trait composition were assessed descriptively using PERMANOVA based on Euclidean distances, and trait-specific group differences were assessed with Kruskal–Wallis tests. Because these groups were defined from the same traits, these tests were interpreted as support for cluster separation rather than as independent ecological tests.

Generalized linear models (GLMs) were used to assess the effects of environmental variables on functional alpha diversity. Functional diversity at q = 0 was used as the response variable because this Hill order gives equal weight to all species and therefore emphasizes rare morphofunctional types. Candidate explanatory variables included canopy cover, litter depth, vegetation strata E1 and E4, and the first two PCA axes derived from vegetation-structure variables (PC1 and PC2). To compare alternative hypotheses about the environmental drivers of functional diversity, we fitted all possible combinations of predictors, including the null model. Model selection was based on Akaike’s Information Criterion (AIC), and models with ΔAIC < 2 were considered plausible. The ten models with the lowest AIC values were retained for comparison. Predictor effects were interpreted from the variables retained in the best-supported models. All statistical analysis and images were created in RStudio.

## 3. Results

### 3.1. Environmental PCA and species composition

The first two components of the PCA on vegetation-structure variables had eigenvalues > 1 and together explained 65.97% of total variance; both axes were retained for subsequent analyses (Fig. 4). The first vegetation PCA axis (PC1) explained 50.71% of the variance and was positively associated with forest age, canopy height and mean DBH, and negatively associated with vegetation stratum E3 (5–10 m) and tree density. This axis represents a gradient of forest maturity and structural complexity: high PC1 scores correspond to older, taller and structurally complex forests, whereas low PC1 scores reflect younger forests with higher tree density at intermediate strata (<10 m). The second axis (PC2) explained 15.26% of the variance and was negatively associated with temperature and intermediate vegetation strata (1–5 m and 5–10 m). Points with high PC2 scores were characterized by smaller trees forming a lower canopy with a pronounced intermediate layer.

**Figure 4.**
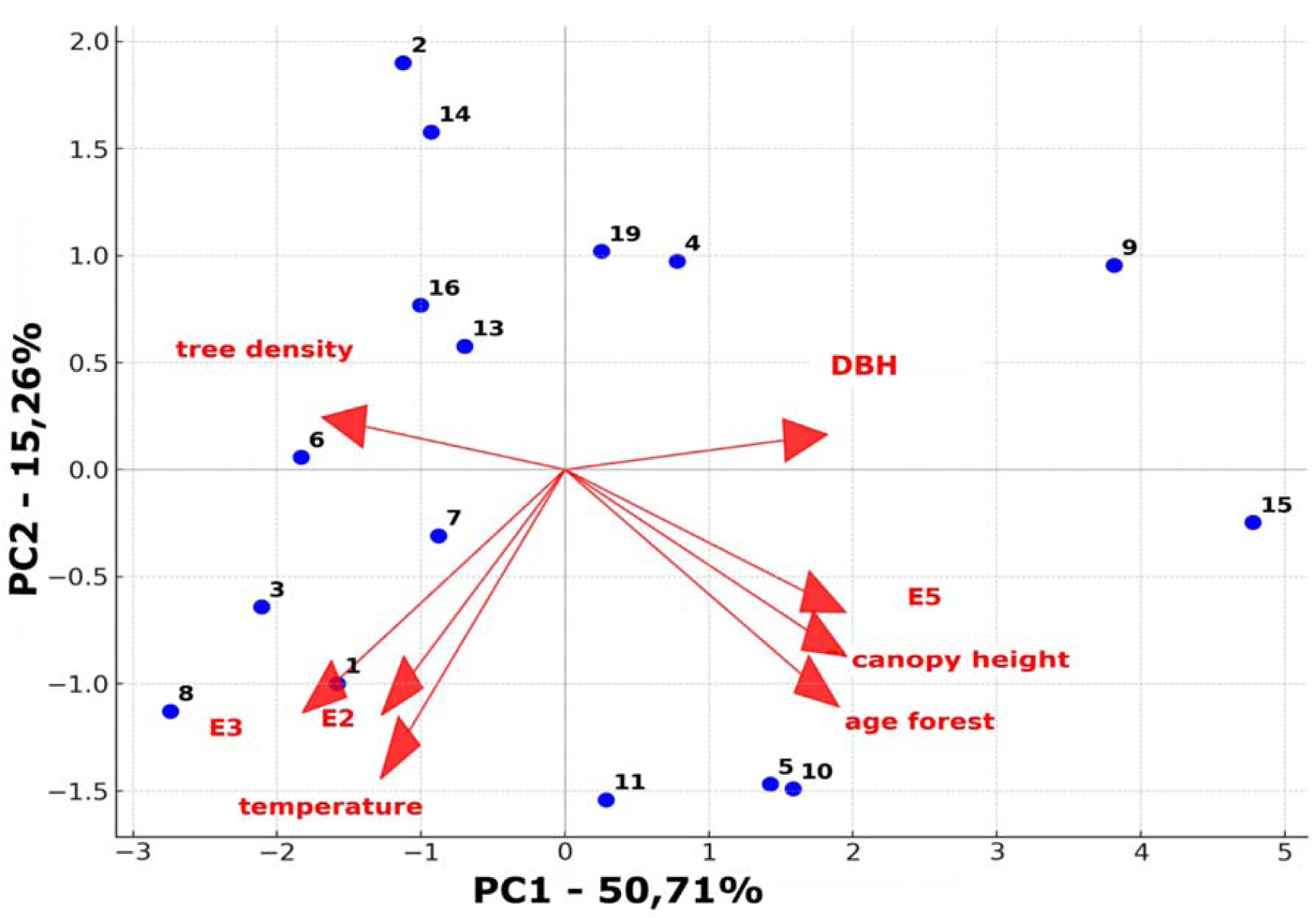
Principal component analysis (PCA) of vegetation-structure variables.. The *X* and *Y* axes represent the two principal components, first principal component (PC1) representing dimension 1 and second principal component (PC2) representing dimension 2, which together explain 65.97% of the total data variation. Blue dots and numbers represent collected areas (see Fig. 1). The variables were abbreviated for analysis purposes, as follows: tree density: tree density per point; DBH: diameter at breast height; canopy height: canopy height; E1: vegetation layer 0–1 m; E2: vegetation layer 1–5 m; E3: vegetation layer 5–10 m; E4: vegetation layer 10–15 m; E5: vegetation layer greater than 15 m; age forest: forest age.

A total of 384 harvestmen were collected, distributed among 14 morphospecies. Gonyleptidae accounted for 57% of the morphospecies collected, followed by Cryptogeobiidae (28%) and Sclerosomatidae (15%). The morphospecies abundance matrix is provided as Supplementary Table A1.

### 3.2. Functional diversity across Hill orders

The Moran’s I analyses revealed no significant spatial autocorrelation for any of the variables examined. Estimates of Moran’s I ranged from weakly negative to moderately positive (–0.20 to 0.32), but all associated p-values were above 0.11, indicating that none of the patterns departed from randomness.

Functional diversity was high across alpha, beta and gamma components, but values depended strongly on the Hill order used to weight species abundance (Fig. 5; Table 1). At all spatial scales, functional diversity was highest at q = 0 and declined as greater weight was assigned to abundant species at q = 1 and q = 2. This pattern indicates that rare species contributed disproportionately to the functional trait space occupied by the harvestman assemblage.

**Figure 5.**
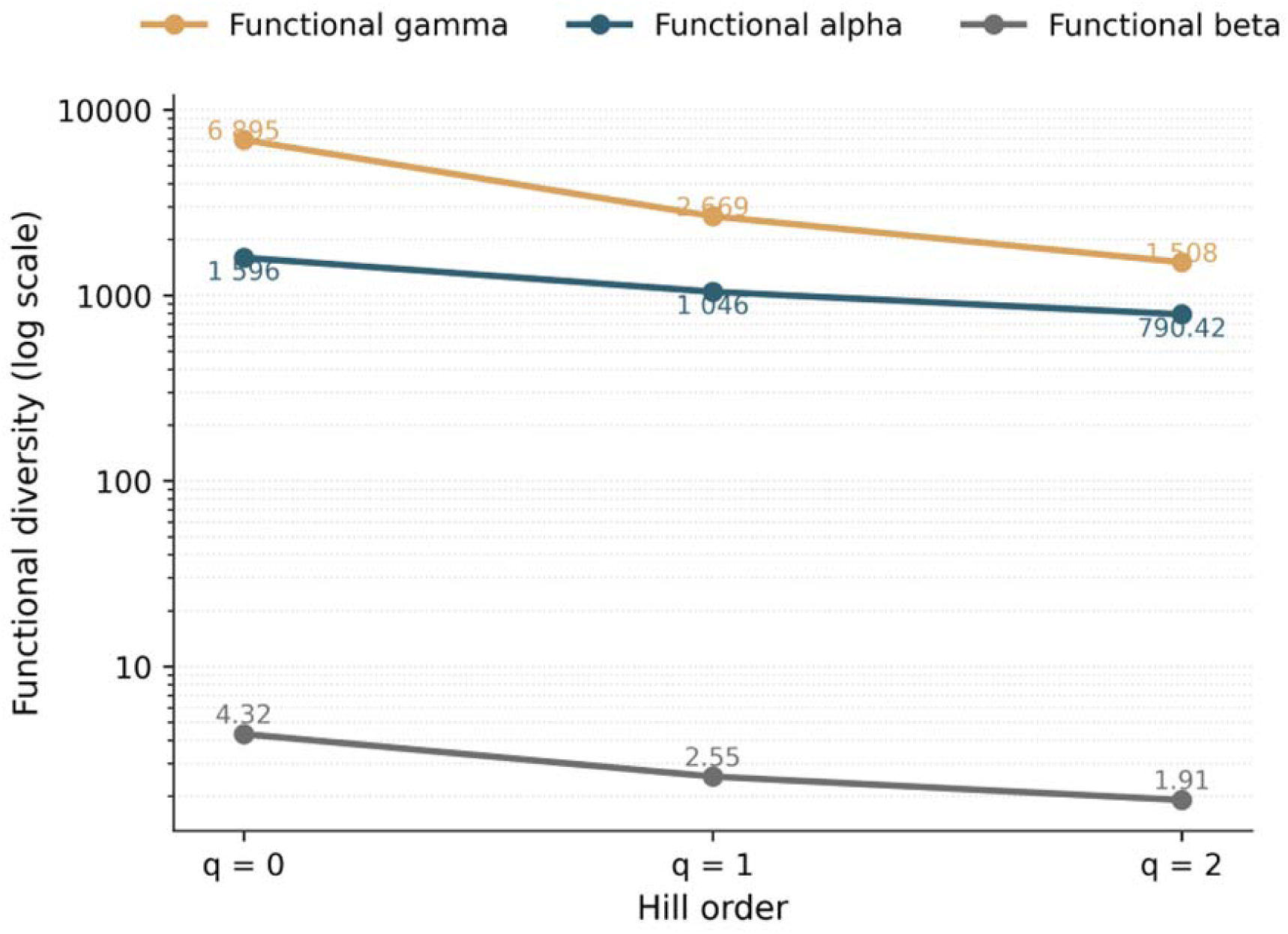
Functional alpha, beta and gamma diversity of harvestmen across Hill orders q = 0, q = 1 and q = 2. The y-axis is shown on a logarithmic scale to allow comparison among diversity components of different magnitudes.

**Table 1.** Functional alpha, beta and gamma diversity of harvestmen based on Hill numbers in Parque Ecológico Imigrantes (PEI), São Paulo, Brazil.

| Hill order | Functional alpha | Functional beta | Functional gamma |
| --- | --- | --- | --- |
| q = 0 | 1595.687 | 4.321 | 6895.123 |
| q = 1 | 1046.495 | 2.550 | 2668.951 |
| q = 2 | 790.421 | 1.908 | 1508.122 |

Functional alpha diversity, representing the average variation in morphological traits within sampling points, decreased from 1595.69 at q = 0 to 790.42 at q = 2. Functional gamma diversity showed an even sharper reduction, from 6895.12 at q = 0 to 1508.12 at q = 2. Thus, dominant species represented only a limited portion of the total functional trait space when compared with the full assemblage including rare species.

### 3.3. Relationship between taxonomic and functional diversity

Taxonomic alpha diversity and functional alpha diversity were strongly and significantly positively correlated across sampling points (Spearman correlation, r= 0,785; p < 0.0001). This relationship suggests that local increases in species number were accompanied by an expansion of morphofunctional trait variation, consistent with functional complementarity among species.

Taxonomic and functional dissimilarities among sampling points were also positively and significantly correlated. The Mantel test comparing Sørensen taxonomic dissimilarity with functional dissimilarity among sampling-point centroids in standardized trait space showed a moderate positive association (Mantel r = 0.532, p < 0.001). A sensitivity analysis based on Bray–Curtis taxonomic dissimilarity and abundance-weighted functional composition produced a similar result (Mantel r = 0.401, p < 0.001). These findings indicate that differences in species composition among sampling points were accompanied by differences in morphofunctional composition.

### 3.4. Partitioning of taxonomic and functional beta diversity

Taxonomic and functional beta diversity showed contrasting partitioning patterns (Table 2; Fig. 6). Mean pairwise taxonomic beta diversity was 0.318, with 43.1% attributed to species turnover (βSIM = 0.137) and 56.9% to nestedness-resultant dissimilarity (βSNE = 0.181).

**Figure 6.**
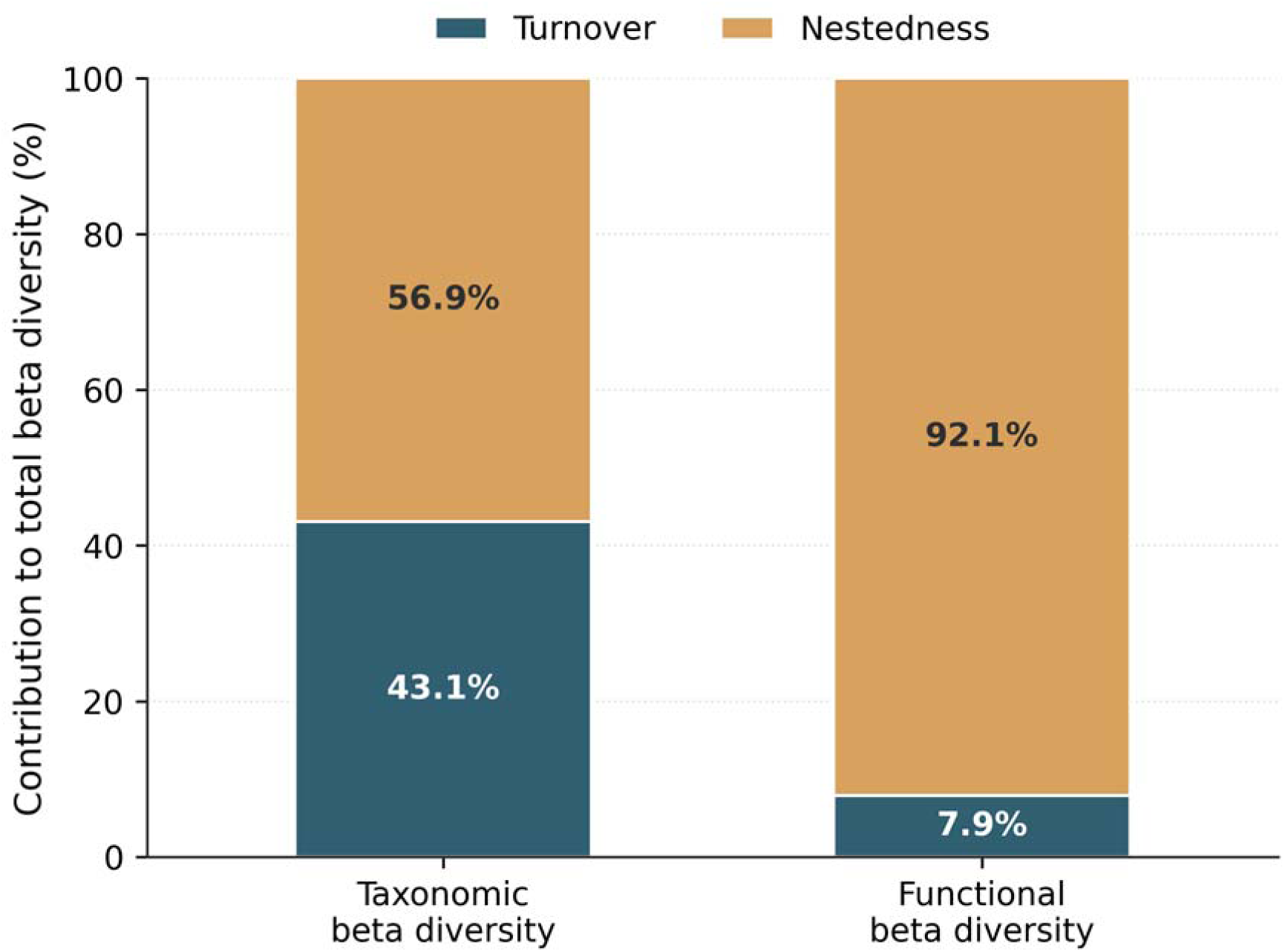
Relative contribution of turnover and nestedness-resultant components to taxonomic and functional beta diversity in harvestman assemblages from Parque Ecológico Imigrantes.

**Table 2.**
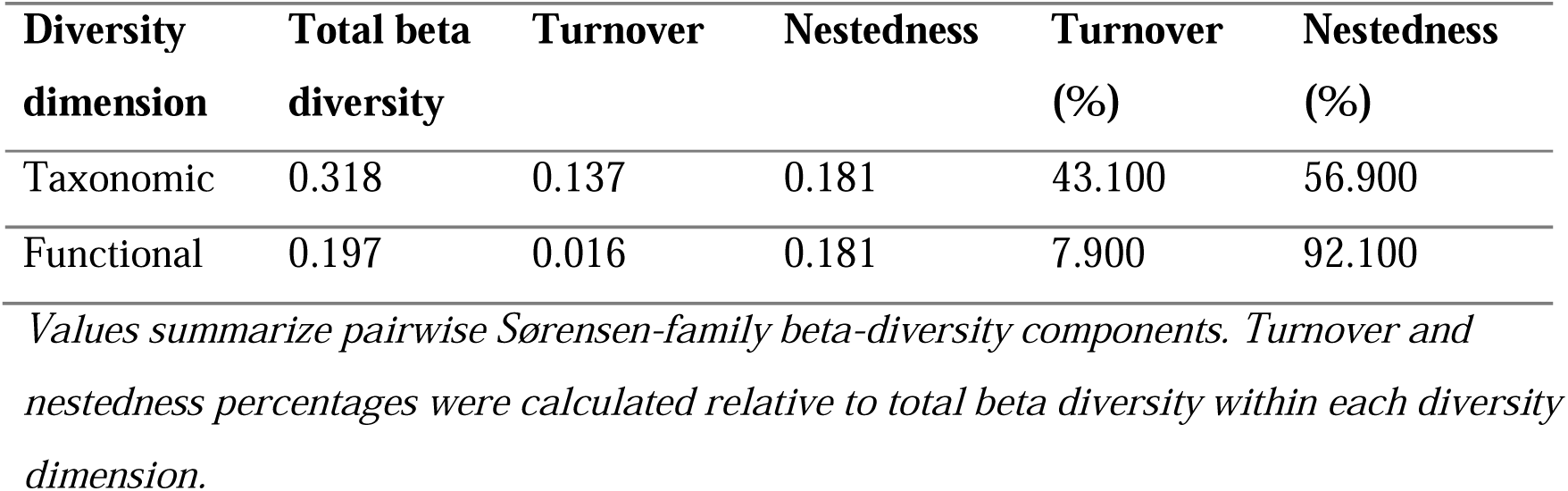
Partitioning of taxonomic and functional beta diversity into turnover and nestedness-resultant components.

| **Diversity dimension** | **Total beta diversity** | **Turnover** | **Nestedness** | **Turnover (%)** | **Nestedness (%)** |
| --- | --- | --- | --- | --- | --- |
| Taxonomic | 0.318 | 0.137 | 0.181 | 43.100 | 56.900 |
| Functional | 0.197 | 0.016 | 0.181 | 7.900 | 92.100 |

*Values summarize pairwise Sørensen-family beta-diversity components. Turnover and nestedness percentages were calculated relative to total beta diversity within each diversity dimension.*

This indicates that differences in species composition among sampling points resulted from both species replacement and the occurrence of species-poor assemblages representing subsets of richer assemblages.

Functional beta diversity was lower than taxonomic beta diversity and was strongly dominated by nestedness-resultant dissimilarity. Total functional beta diversity was 0.197, of which only 7.9% corresponded to functional turnover (0.016), whereas 92.1% corresponded to functional nestedness (0.181) (Table 2). Thus, differences in functional composition among sampling points were driven mainly by variation in the amount of functional space occupied by local assemblages, rather than by the replacement of distinct functional strategies among sites.

Mantel tests showed significant positive associations between taxonomic and functional dissimilarity components (Table 3). Total taxonomic and functional beta diversity were positively correlated (Mantel r = 0.659, p < 0.001). Significant positive relationships were also found between taxonomic and functional turnover (Mantel r = 0.521, p < 0.001) and between taxonomic and functional nestedness-resultant components (Mantel r = 0.636, p < 0.001). These results indicate that spatial changes in species composition were accompanied by changes in functional composition, although the dominant functional pattern was nested loss or contraction of trait space across sampling points.

**Table 3.** Mantel tests between taxonomic and functional beta-diversity components.

| **Comparison** | **Mantel r** | **P-value** | **Interpretation** |
| --- | --- | --- | --- |
| Total taxonomic beta × total functional beta | 0.659 | <0.001 | Sites differing in species composition also differed functionally. |
| Taxonomic turnover × functional turnover | 0.521 | <0.001 | Species replacement was associated with some functional replacement. |
| Taxonomic nestedness × functional nestedness | 0.636 | <0.001 | Species-poor subsets were associated with functional-space contraction. |

*Mantel tests were based on pairwise dissimilarity matrices and 9,999 permutations.*

### 3.5. Morphofunctional structure and trait contributions

The PCA of morphofunctional traits showed that PC1 explained 71.76% of total trait variation and PC2 explained 20.73%, together summarizing 92.49% of the morphofunctional space (Fig. 7; Table 4). PC1 was positively correlated with all four traits, especially body length, body width, femur IV length and leg II length, and therefore represented an overall size and appendage-length gradient. PC2 contrasted appendage elongation against body robustness: leg II and femur IV loaded positively, whereas body length and body width loaded negatively.

**Figure 7.**
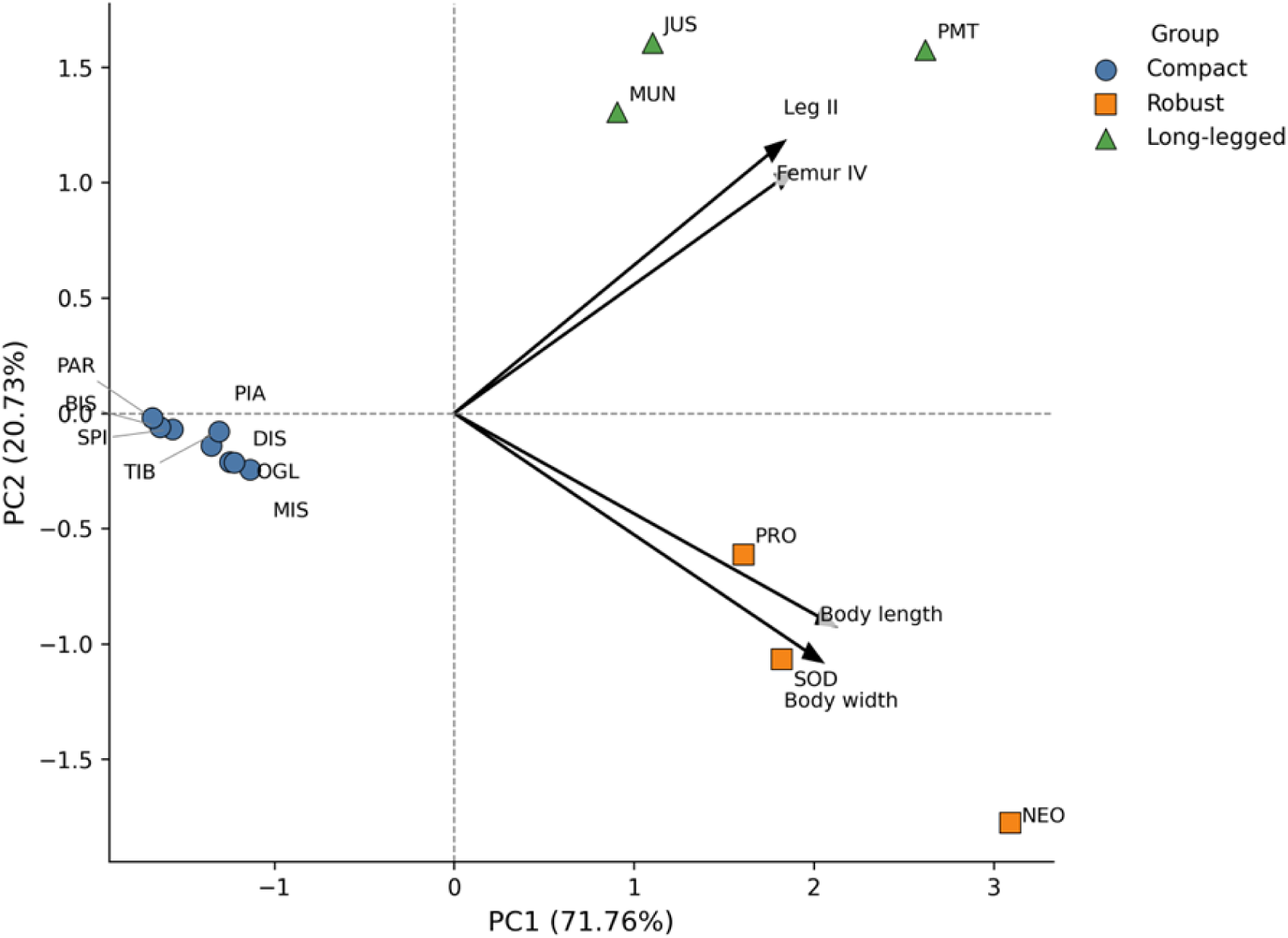
Principal component analysis (PCA) of harvestman morphofunctional traits based on body length, body width, femur IV length and leg II length. Species are plotted according to their scores on the first two PCA axes, and arrows represent the direction and magnitude of trait contributions. Colors/symbols indicate the three morphofunctional groups identified in the clustering analysis: compact, robust and long-legged species. Species codes: JUS = Jussara sp.; MUN = Munequita pulchra; DIS = Discocyrtus rarus; MIS = Mischonyx squalidus; NEO = Neosadocus maximus; OGL = Ogloblinia loretoensis; PIA = Piassagera brieni; PRO = Progonyleptoidellus striatus; PMT = Promitobates ornatus; SOD = Sodreana sodreana; TIB = Tibangara sp.; SPI = Spinopilar sp.; BIS = Bissula paradoxa; PAR = Paratricommatus sp.

**Table 4.** Trait loadings and correlations with the first two morphofunctional PCA axes.

| **Trait** | **PC1 loading** | **PC2 loading** | **r with PC1** | **p with PC1** | **r with PC2** | **p with PC2** |
| --- | --- | --- | --- | --- | --- | --- |
| Body length | 0.536 | -0.434 | 0.908 | <0.001 | -0.395 | 0.162 |
| Body width | 0.518 | -0.507 | 0.877 | <0.001 | -0.462 | 0.096 |
| Femur IV | 0.479 | 0.497 | 0.811 | <0.001 | 0.452 | 0.104 |
| Leg II | 0.464 | 0.555 | 0.787 | <0.001 | 0.505 | 0.065 |

Hierarchical clustering identified three morphofunctional groups: compact species, robust large-bodied species and long-legged species (Fig. 7). Compact species had small bodies and short appendages; robust species had larger and wider bodies; and long-legged species had proportionally elongated appendages. Group separation was strong in the standardized trait space (PERMANOVA: F = 32.14, R² = 0.854, p < 0.001), and the groups differed in each of the four individual traits (Kruskal–Wallis tests: H = 10.37, p = 0.0056 for body length, body width, femur IV length and leg II length).

### 3.6. Environmental drivers of functional alpha diversity

Model selection indicated that functional alpha diversity at q = 0 was best explained by canopy cover, litter depth and the forest-maturity gradient represented by PC1. The best-supported model included canopy cover, litter depth and PC1 (AIC = 121.42, ΔAIC = 0), whereas the second-best model additionally included PC2 and was also considered plausible (AIC = 123.32, ΔAIC = 1.90; Fig.8; Table 5). Models including E1 or E4 received weaker support, with ΔAIC values greater than 2. The null model had very low support (AIC = 141.58, ΔAIC = 20.16), indicating that vegetation structure substantially improved the explanation of variation in functional alpha diversity. Overall, these results suggest that canopy structure, litter depth and forest maturity were the most important predictors of harvestman functional diversity.

**Figure 8.**
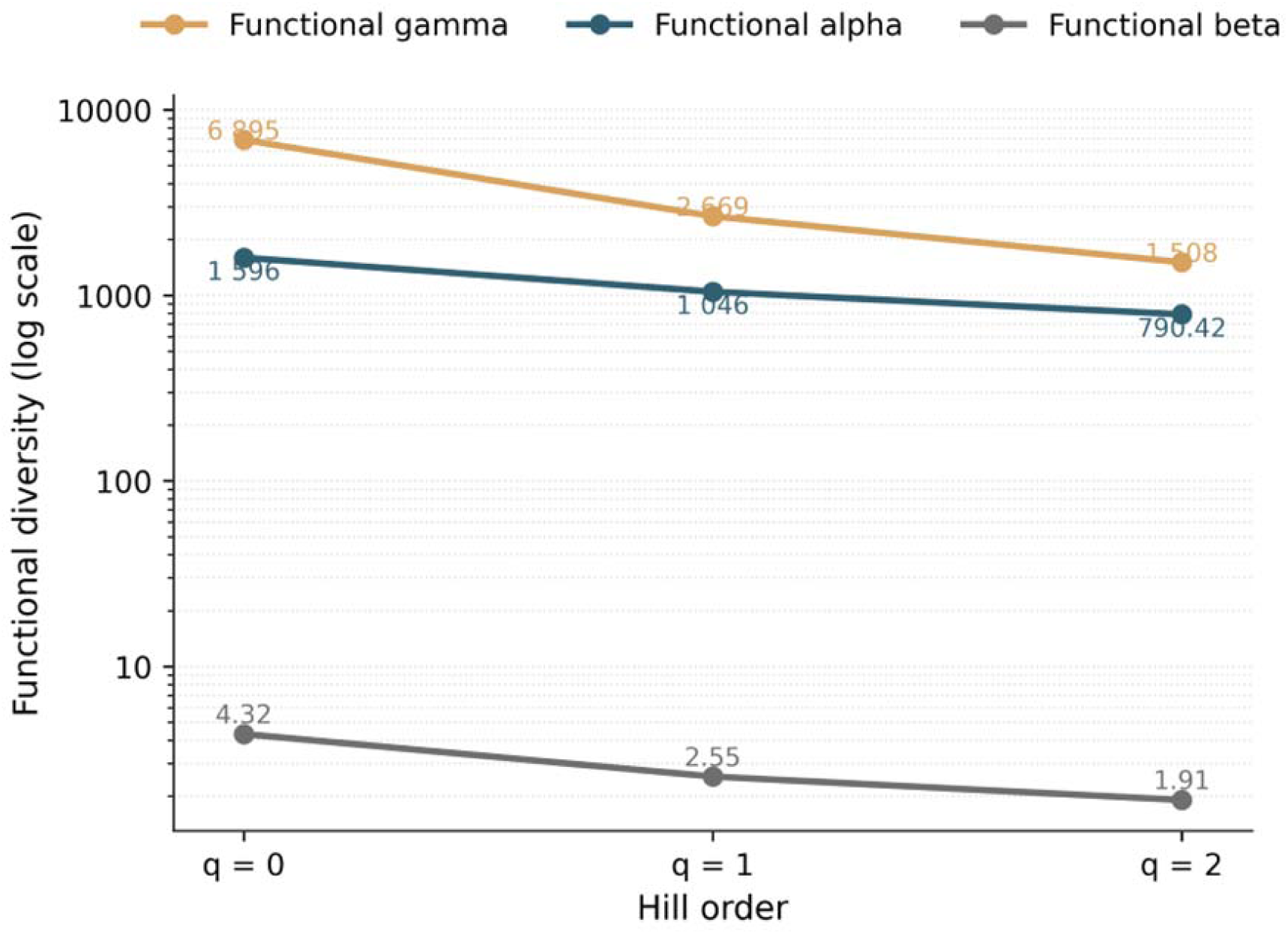
AIC-based selection of environmental predictors of harvestman functional alpha diversity. The left panel shows ΔAIC values for candidate models, with the dashed vertical line indicating the ΔAIC = 2 threshold. The right panel indicates which predictors were included in each model. Highlighted rows correspond to plausible models.

**Table 5.** Model selection based on Akaike’s Information Criterion for generalized linear models explaining harvestman functional alpha diversity at q = 0.

| Rank | Model | Predictors | AIC | $\Delta AIC$ | Support |
| --- | --- | --- | --- | --- | --- |
| 1 | M1 | Canopy cover + litter depth + PC1 | 121.42 | 0.00 | Plausible |
| 2 | M2 | Canopy cover + litter depth + PC1 + PC2 | 123.32 | 1.90 | Plausible |
| 3 | M3 | Canopy cover + E1 + E4 + litter depth | 123.71 | 2.29 | Lower support |
| 4 | M4 | Canopy cover + E1 + PC2 | 124.00 | 2.58 | Lower support |
| 5 | M5 | Canopy cover + E1 + E4 + litter depth + PC1 | 125.40 | 3.97 | Lower support |
| 6 | M6 | Canopy cover + E1 + E4 + PC1 + PC2 | 125.94 | 4.52 | Lower support |
| 7 | M7 | Canopy cover + E1 + PC1 | 129.09 | 7.67 | Lower support |
| 8 | M8 | Canopy cover + E4 + litter depth | 130.15 | 8.73 | Lower support |
| 9 | M9 | Canopy cover + E4 + litter depth + PC2 | 130.38 | 8.95 | Lower support |
| 10 | M10 | Canopy cover + E4 + litter depth + PC1 | 130.78 | 9.35 | Lower support |
| — | Null | Intercept only | 141.58 | 20.16 | Null model |

**Note.** Models are ranked by increasing AIC. Models with $\Delta AIC < 2$ were considered plausible. PC1 and PC2 correspond to the first two axes of the vegetation-structure PCA. The null model corresponds to the intercept-only model.

## 4. Discussion

This study provides one of the first functional-trait perspectives on harvestman communities in the Atlantic Forest and shows that forest structure, rare species and functional-space nestedness are central to the organization of local assemblages. The main pattern was consistent across diversity components: functional diversity was highest when rare species were weighted equally and declined when abundant species received greater influence. This indicates low functional redundancy and suggests that uncommon harvestmen carry a substantial portion of trait variation.

The positive relationship between taxonomic and functional alpha diversity indicates that species-rich sampling points also tended to contain a broader range of morphologies. This pattern supports the interpretation that species accumulation in these communities is not merely numerical but also adds functional complementarity [20]. For conservation, this is important because the loss of rare or locally restricted species could translate into the loss of unique combinations of body size, body robustness and appendage length, reducing the functional breadth of the assemblage [18,22].

The Mantel analyses further support the link between taxonomic and functional dimensions of diversity. Sampling points that differed more strongly in species composition also differed more strongly in morphofunctional composition, indicating that taxonomic dissimilarity was not merely associated with the replacement of functionally equivalent species. Similar approaches have been used in trait-based studies to evaluate whether changes in taxonomic composition are accompanied by changes in functional composition [26,5]. For example, Stefanidis [39] found significant correlations between taxonomic and functional beta diversity in aquatic macrophyte communities, reinforcing the value of comparing both dimensions to understand community responses to environmental gradients.

The formal decomposition of beta diversity refines the interpretation of spatial variation in harvestman assemblages. Taxonomic beta diversity was explained by both turnover and nestedness, indicating that sampling points differed partly because of species replacement and partly because some local assemblages contained subsets of the species present in richer assemblages. In contrast, functional beta diversity was overwhelmingly dominated by nestedness. This suggests that differences among sampling points did not primarily reflect the replacement of one set of morphofunctional strategies by another, but rather variation in how much of the regional functional space was represented locally.

This result changes the ecological interpretation of functional beta diversity in an important way. Although rare species contributed substantially to regional functional diversity, among-site functional differentiation appears to be structured mainly by functional-space contraction in some assemblages. In other words, sites with lower functional diversity seem to retain only a subset of the morphofunctional strategies present in richer or more structurally developed sites. Such a pattern is consistent with low functional redundancy: the loss of species, especially rare or morphologically distinctive ones, may reduce the amount of functional space occupied locally even when communities are not replaced by completely different functional groups.

The predominance of functional nestedness also strengthens the conservation interpretation of the study. If younger or structurally simplified forest patches contain only subsets of the functional strategies found in older and more complex forest patches, then mature forest structure may be necessary to maintain the full regional spectrum of harvestman morphologies. This pattern differs from studies in which functional beta diversity is mainly driven by turnover, where distinct habitats harbor different functional strategies [23,12,44]. Here, the strong nestedness component suggests a process closer to functional impoverishment or filtering, in which some sites lose portions of the regional functional space.

The ordination analyses identified three broad morphofunctional groups: compact species, robust large-bodied species and long-legged species. Although the present study did not directly quantify microhabitat use for each species, these morphologies are ecologically interpretable. Compact species may be favored in leaf litter and narrow refuges, robust species may use exposed surfaces such as trunks, leaf or broader substrates, and long-legged forms may be associated with movement on vegetation or elevated structures. These interpretations are consistent with the known sensitivity of harvestmen to moisture, substrate structure and forest disturbance [21,30,3], but future field observations are necessary to validate the behavioral and ecological meaning of these morphofunctional groups.

Environmental model selection showed that functional alpha diversity was most strongly associated with canopy cover, litter depth and the vegetation-structure gradient represented by PC1. This result reinforces the importance of both near-ground habitat structure and broader forest maturity for maintaining morphofunctionally diverse harvestman assemblages. Deeper litter may increase the availability of refuges, foraging substrates and humid microhabitats, whereas canopy cover and mature forest structure may contribute to greater microclimatic stability [17,35]. Although PC2 appeared in a second plausible model, its support was weaker than that of canopy cover, litter depth and PC1. Vegetation strata E1 and E4 were not retained in the best-supported models, suggesting that fine-scale vertical vegetation layers may be less important than litter accumulation, canopy conditions and overall forest maturity in explaining functional alpha diversity at this local scale.

These results strengthen the view that mature Atlantic Forest remnants are disproportionately important for preserving the functional dimension of arthropod biodiversity [3,40,28].Although secondary forests can recover species richness over time, functional recovery may depend on the re-establishment of specific habitat structures, including litter accumulation, canopy development and vertical complexity. In this study, patches older than 60 years had the highest functional alpha diversity, indicating that long-term structural regeneration may be necessary for maintaining rare morphofunctional strategies. This has direct implications for conservation planning in the Atlantic Forest, where remaining forest is often embedded in fragmented and regenerating landscapes [36,47].

Taken together, our results show that conservation strategies should not only aim to maximize species richness, but also to prevent the local loss of rare and functionally distinctive morphologies that contribute to the functional breadth of Atlantic Forest harvestman assemblages. The nestedness-dominated pattern found here suggests that functionally simplified sites may not be equivalent alternatives to mature sites, because they may conserve only partial subsets of the regional trait space.

## 5. Conclusions

Harvestman communities in Parque Ecológico Imigrantes showed high functional diversity, low functional redundancy and a strong contribution of rare species to regional morphofunctional variation. Functional alpha diversity was positively associated with taxonomic alpha diversity, indicating that species-rich points also contained broader morphological variation. However, beta-diversity partitioning showed that functional differences among sampling points were dominated by nestedness rather than turnover, revealing a pattern of functional-space contraction in poorer assemblages.

Forest structural maturity, canopy cover and litter depth were the main environmental predictors of functional diversity, with older and structurally more developed forest patches supporting a more complete representation of the regional functional trait space. These findings suggest that the recovery and conservation of mature forest structure are essential for maintaining functionally complete harvestman assemblages in the Atlantic Forest. More broadly, the study highlights harvestmen as promising models for functional ecology in tropical forests and opens new avenues for linking morphology, microhabitat use and conservation value in Neotropical arachnids.

## Acknowledgements

We are grateful to Parque Ecológico Imigrantes for authorizing and facilitating the field surveys, for providing field guides during collection activities, and for supplying logistical support and sampling materials essential to this study. To Juliana Abud and Luana Camargo for their assistance with vegetation structure data, Thomas Puttker for the sampling design and Fabiana Casarin for their assistance with field collections

## Credit authorship contribution statement

R.C.C.: Investigation, Data curation, Formal analysis, Writing, original draft

L.S.L: Investigation, Data curation, Formal Analysis

B.K.S: Formal analysis, Writing, review and editing

S.R.: Formal analysis, Writing, review and editing

C.B.: Conceptualization, Methodology, Supervision, Project administration, Resources, Writing, review and editing.

## Data availability statement

Data are available from the corresponding author upon reasonable request.

## Supplementary material

**Supplementary Table A1.**
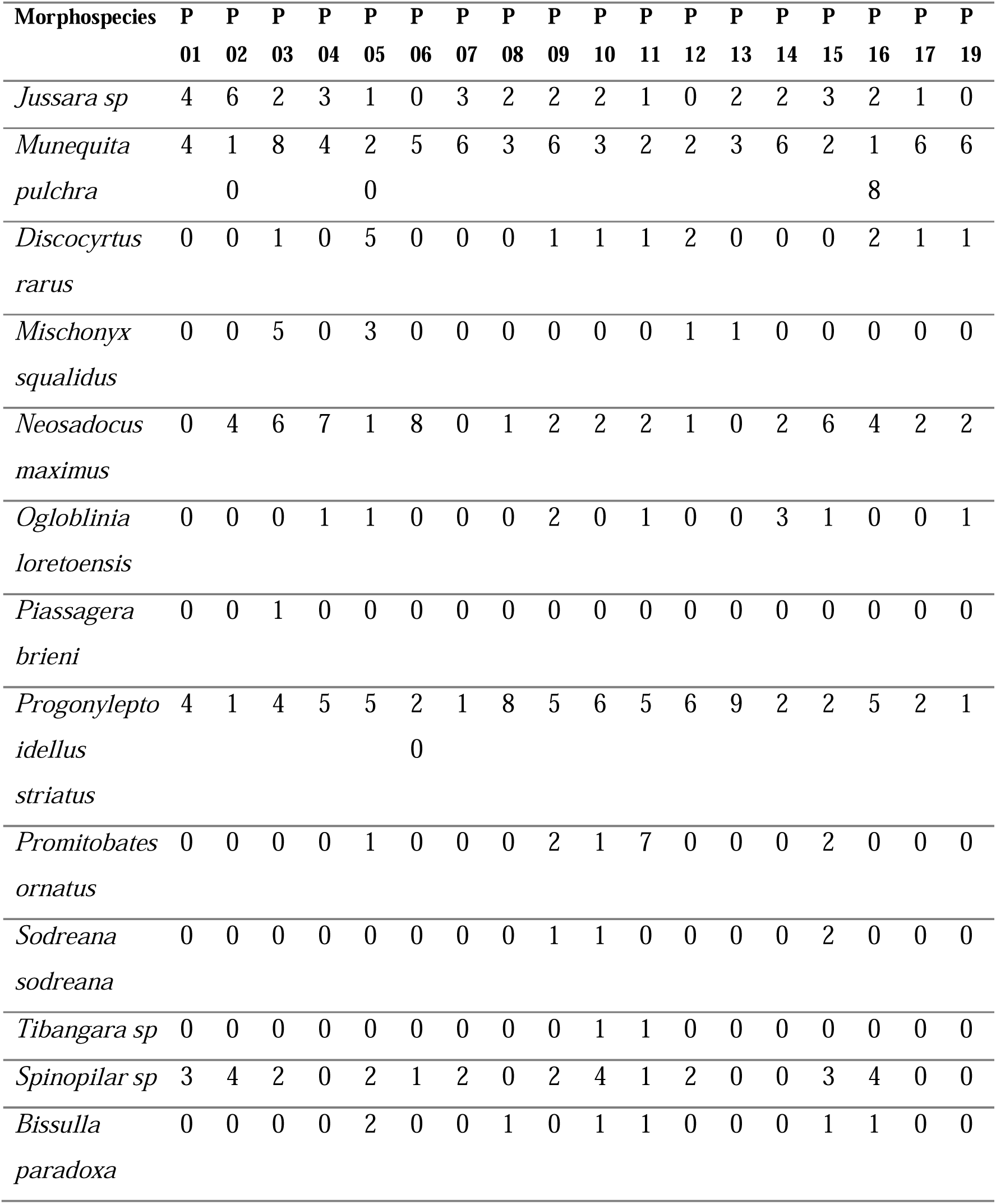
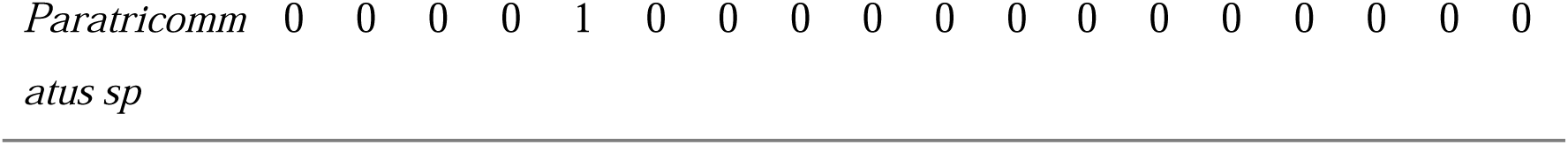
Morphospecies abundance matrix of harvestmen collected in Parque Ecológico Imigrantes, São Paulo, Brazil.

**Supplementary Table A2.** Number of individuals measure of each morphospecies.

| Morphospecies | Male | Female | Indeterminate |
| --- | --- | --- | --- |
| *Jussara sp* | 5 | 0 | - |
| *Munequita pulchra* | 5 | 0 | - |
| *Discocyrtus rarus* | 9 | 4 | - |
| *Mischonyx squalidus* | 4 | 3 | - |
| *Neosadocus maximus* | 10 | 10 | - |
| *Ogloblinia loretoensis* | 0 | 12 | - |
| *Piassagera brieni* | 1 | 0 | - |
| *Progonyleptoidellus striatus* | 6 | 14 | - |
| *Promitobates ornatus* | 3 | 2 | - |
| *Sodreana sodreana* | 0 | 2 | - |
| *Tibangara sp* * | 0 | 0 | 1 |
| *Spinopilar sp* | 10 | 10 | - |
| *Bissulla paradoxa* * | 0 | 0 | 2 |
| *Paratricommatus sp* * | 0 | 0 | 1 |

\*The sex of the individuals in the morphospecies *Tibangara sp*, *Bissulla paradoxa* and *Paratricommatus sp* was inconclusive.

## References

[1] Baselga A. Partitioning the turnover and nestedness components of beta diversity. Global Ecology and Biogeography, 2010, 19: 134.143. doi: 10.1111/j.1466-8238.2009.00490.x.

[2] Baselga A, Orme C D L. betapart: an R package for the study of beta diversity. Methods in Ecology and Evolution, 2012, 3: 808.812. doi: 10.1111/j.2041-210X.2012.00224.x.

[3] Bragagnolo C, Pinto-da-Rocha R. Harvestmen in an Atlantic forest fragmented landscape: evaluating assemblage response to habitat quality and quantity. Biological Conservation, 2007, 139 (3-4): 389.400. doi: 10.1016/j.biocon.2007.07.008.

[4] Bueno A S, et al. Post-disturbance recovery of functional diversity in tropical forest communities. Journal of Applied Ecology, 2022, 59: 1434.1445.

[5] Camargo-Vanegas J, de la Hoz-Pedraza S, Sierra-Chamorro H, Guerrero R J. The Taxonomic and Functional Diversity of Leaf-Litter Dwelling Ants in the Tropical Dry Forest of the Colombian Caribbean. Diversity, 2024, 16 (11): 687. doi: 10.3390/d16110687.

[6] Chase J, McGill B, Thompson P, Antão L, Bates A, Blowes S, Dornelas M, Gonzalez A, Magurran A, Supp S, et al. Species richness change across spatial scales. Oikos, 2019, 128 (8): 1079.1091. doi: 10.1111/oik.05968.

[7] Chiu C, Chao A. Distance-Based Functional Diversity Measures and Their Decomposition: A Framework Based on Hill Numbers. PLOS One, 2014, 9 (11): e100014. doi: 10.1371/journal.pone.0100014.

[8] CLIMATE-DATA.ORG. [cited 2022 Feb 10]. Available from: https://en.climate-data.org/

[9] DaSilva M B, Pinto-da-Rocha R. A história biogeográfica da Mata Atlântica: opiliões (Arachnida) como modelo para sua inferência. In: Carvalho C J B, Almeida E A B, eds., Biogeografia Da América Do Sul - Padrões E Processos, São Paulo, Editora Roca, 2011. 221.238.

[10] Dornelas M, Chase J, Gotelli N, Magurran A, McGill B, Antão L, Blowes S, Daskalova G, Leung B, Martins I, Moyes F, Myers-Smith I, Thomas C, Vellend M. Looking back on biodiversity change: lessons for the road ahead. Philosophical Transactions of the Royal Society B: Biological Sciences, 2023, 378 (1881): 20220199. doi: 10.1098/rstb.2022.0199.

[11] Flynn D F B, Gogol-Prokurat M, Nogeire T, Molinari N, Richers B T, Lin B B, Simpson N, Mayfield M M, DeClerck F. Loss of functional diversity under land use intensification across multiple taxa. Ecology Letters, 2009, 12 (1): 22.33. doi: 10.1111/j.1461-0248.2008.01255.x.

[12] Hernández-Mendoza L C, Escalera-Vázquez L H, Vega-Cendejas M E, Núñez-Lara E, Chiappa-Carrara X, Arceo-Carranza D. Functional and taxonomic β diversity in fish assemblages is structured by turnover in a tropical coastal lagoon. Environmental Biology of Fishes, 2024, 107 (6): 1219.1234. doi: 10.1007/s10641-024-01626-y.

[13] Higgins K F, Jenkins K J, Clambey G K, Uresk D W, Naugle D E. Vegetation Sampling and Measurement. In: Iowa State University, From the SelectedWorks of Robert Klaver, Ames, 2012. 381.409.

[14] Hillebrand H, Blasius B, Borer E T, Chase J M, Downing J A, Eriksson B K, Filstrup C T, Harpole W S, Hodapp D, Larsen S, Lewandowska A M, Seabloom E W, Van de Waal D B, Ryabov A B. Biodiversity change is uncoupled from species richness trends: Consequences for conservation and monitoring. Journal of Applied Ecology, 2018, 55 (1): 169.184. doi: 10.1111/1365-2664.12959.

[15] Koleff P, Gaston K J, Lennon J J. Measuring beta diversity for presence–absence data. Journal of Animal Ecology, 2003, 72 (3): 367.382. doi: 10.1046/j.1365-2656.2003.00710.x.

[16] Kury A B, Mendes A C, Cardoso L, Kury M S, Granado A A. WCO-Lite: online world catalogue of harvestmen (Arachnida, Opiliones). Version 1.0 – Checklist of all valid nomina in Opiliones with authors and dates of publication up to 2018. Rio de Janeiro: The Authors, 2020. ISBN 978-65-00-06706-4.

[17] Lian Z, Fan C, Wang J, von Gadow K. Structure complexity is the primary driver of functional diversity in the temperate forests of northeastern China. Forest Ecosystems, 2022, 9: 100048. doi: 10.1016/j.fecs.2022.100048.

[18] Leitão R P, Zuanon J, Villéger S, Williams S E, Baraloto C, Fortunel C, Mendonça F P, Mouillot D. Rare species contribute disproportionately to the functional structure of species assemblages. Proceedings of the Royal Society B: Biological Sciences, 2016, 283 (1828): 20160084. doi: 10.1098/rspb.2016.0084.

[19] Magurran A E. Measuring biological diversity. Oxford, Blackwell Publishing, 2004. ISBN 978-0-632-05633-0.

[20] Maherali H, Klironomos J N. Influence of Phylogeny on Fungal Community Assembly and Ecosystem Functioning. Science, 2007, 316 (5832): 1746.1748. doi: 10.1126/science.1143082.

[21] Mestre L A M, Pinto-da-Rocha R. Population dynamics of an isolated population of the harvestman Ilhaia cuspidata (Opiliones, Gonyleptidae) in Araucaria Forest (Curitiba, Paraná, Brazil). The Journal of Arachnology, 2004, 32 (2): 208.220. doi: 10.1636/m02-61.

[22] Mouillot D, Graham N A J, Villéger S, Mason N W H, Bellwood D R. A functional approach reveals community responses to disturbances. Trends in Ecology & Evolution, 2013, 28 (3): 167.177. doi: 10.1016/j.tree.2012.10.004.

[23] Myers E M V, Eme D, Liggins L, Harvey E S, Roberts C D, Anderson M J. Functional beta diversity of New Zealand fishes: Characterising morphological turnover along depth and latitude gradients, with derivation of functional bioregions. Austral Ecology, 2021, 46 (6): 965.981. doi: 10.1111/aec.13078.

[24] Nogueira A A, Bragagnolo C, DaSilva M B, Martins T K, Lorenzo E P, Perbiche-Neves G, Pinto-da-Rocha R. Historical signatures in the alpha and beta diversity patterns of Atlantic

[25] Forest harvestman communities (Arachnida: Opiliones). Canadian Journal of Zoology, 2019, 97 (7): 631.643. doi: 10.1139/cjz-2018-0032.

[26] Núñez J C, Romero M A, Reinaldo M O, González R, Magurran A, Svendsen G. Species turnover drives functional turnover with balanced functional richness in a Patagonian demersal assemblage. Journal of Sea Research, 2023, 195: 102452. doi: 10.1016/j.seares.2023.102452.

[27] Pardini R, de Souza S M, Braga-Neto R, Metzger J P. The role of forest structure, fragment size and corridors in maintaining small mammal abundance and diversity in an Atlantic forest landscape. Biological Conservation, 2005, 124 (2): 253.266. doi: 10.1016/j.biocon.2005.01.033.

[28] Pardini R, Faria D, Accacio G M, Laps R R, Mariano-Neto E, Paciencia M L B, Dixo M, Baumgarten J. The challenge of maintaining Atlantic forest biodiversity: A multi-taxa conservation assessment of specialist and generalist species in an agro-forestry mosaic in southern Bahia. Biological Conservation, 2009, 142 (6): 1178.1190. doi: 10.1016/j.biocon.2009.02.010.

[29] Petchey O L, Gaston K J. Functional diversity: back to basics and looking forward. Ecology Letters, 2006, 9 (6): 741.758. doi: 10.1111/j.1461-0248.2006.00924.x.

[30] Pinto-da-Rocha R, Machado G, Giribet G, eds. Harvestmen: The Biology of Opiliones. Cambridge, Harvard University Press, 2007. ISBN 9780674023437.

[31] Pinto-da-Rocha R, DaSilva M B, Bragagnolo C. The faunistic similarity and historic biogeography of the harvestmen of southern and southeastern Atlantic rain forest of Brazil. The Journal of Arachnology, 2005, 33 (2): 290.299. doi: 10.1636/04-114.1.

[32] Proud D N, Felgenhauer B E, Townsend Jr V R, Osula D O, Gilmore III W O, Napier Z L, Van Zandt P A. Diversity and habitat use of Neotropical harvestmen (Arachnida: Opiliones) in a Costa Rican rainforest. International Scholarly Research Notices, 2012, 2012 (1): 549765. doi: 10.5402/2012/549765.

[33] Raub F, Höfer H, Scheuermann L, de Britez R M, Brandl R. Conserving landscape structure – conclusions from partitioning of spider diversity in southern Atlantic forests of Brazil. Studies on Neotropical Fauna and Environment, 2015, 50 (3): 158.174. doi: 10.1080/01650521.2015.1071959.

[34] Rodrigues A C, Villa P M, Ali A, Ferreira-Júnior W, Neri A V. Fine-scale habitat differentiation shapes the composition, structure and aboveground biomass but not species richness of a tropical Atlantic forest. Journal of Forestry Research, 2020, 31 (5): 1599.1611. doi: 10.1007/s11676-019-00994-x.

[35] Romoth K, Darr A, Papenmeier S, Zettler M, Gogina M. Substrate Heterogeneity as a Trigger for Species Diversity in Marine Benthic Assemblages. Biology, 2023, 12 (6): 825. doi: 10.3390/biology12060825.

[36] Rozendaal D M A, Bongers F, Aide T M, Alvarez-Davila E, Ascarrunz N, Balvanera P, Becknell J M, Bentos T V, Brancalion P H S, Cabral G A L, et al. Biodiversity recovery of Neotropical secondary forests. Science Advances, 2019, 5 (3): eaau3114. doi: 10.1126/sciadv.aau3114.

[37] Schneider C A, Rasband W S, Eliceiri K W. NIH Image to ImageJ: 25 years of image analysis. Nature Methods, 2012, 9 (7): 671.675. doi: 10.1038/nmeth.2089.

[38] Schowalter T D, Noriega J A, Tscharntke T. Insect effects on ecosystem services—Introduction. Basic and Applied Ecology, 2018, 26 (6): 1.7. doi: 10.1016/j.baae.2017.09.011.

[39] Stefanidis K, Oikonomou A, Dimitrellos G, Tsoukalas D, Papastergiadou E. Beyond taxonomic diversity patterns – investigating how α and β components of macrophyte functional diversity respond to environmental gradients in lotic ecosystems of Greece. Frontiers in Plant Science, 2023, 14: 1204383. doi: 10.3389/fpls.2023.1204383.

[40] Tsafack N, Fattorini S, Boieiro M, Rigal F, Ros-Prieto A, Ferreira M T, Borges P A V. The Role of Small Lowland Patches of Exotic Forests as Refuges of Rare Endemic Azorean Arthropods. Diversity, 2021, 13 (9): 443. doi: 10.3390/d13090443.

[41] Tourinho A L M. 2007. Padrões de distribuição e fatores condicionantes da riqueza e composição de opiliões na várzea do Rio Amazonas, Brasil (Arachnida: Opiliones). Dissertation. Instituto Nacional de Pesquisas da Amazônia (INPA), Manaus.

[42] Villéger S, Grenouillet G, Brosse S. Decomposing functional beta-diversity reveals that low functional beta-diversity is driven by low functional turnover in European fish assemblages. Global Ecology and Biogeography, 2013, 22 (6): 671.681. doi: 10.1111/geb.12021.

[43] Wang H, Fu H, Wen Z, Yuan C, Zhang X, Ni L, Cao T. Seasonal patterns of taxonomic and functional beta diversity in submerged macrophytes at a fine scale. Ecology and Evolution, 2021, 11 (14): 9827.9836. doi: 10.1002/ece3.7811.

[44] Wang J, Jiang S, Sun D, Chen J, Li B, Chen L. Turnover structures in macrobenthic communities rather than nestedness in the Yellow River Delta wetland, China. Aquatic Sciences, 2024, 86 (4): 99. doi: 10.1007/s00027-024-01118-2.

[45] Ward J H Jr. Hierarchical grouping to optimize an objective function. Journal of the American Statistical Association, 1963, 58 (301): 236.244. doi: 10.1080/01621459.1963.10500845.

[46] Webb C O, Ackerly D D, McPeek M A, Donoghue M J. Phylogenies and community ecology. Annual Review of Ecology and Systematics, 2002, 33: 475.505. doi: 10.1146/annurev.ecolsys.33.010802.150448.

[47] Wei J, Chen H, Yu X, Guo Z, Zhang X, Tian L, Nong S. Stand structure and plant diversity characteristics of typical artificial forests after natural recovery in the hilly region of central Hainan. Frontiers in Plant Science, 2025, 16: 1629250. doi: 10.3389/fpls.2025.1629250.

[48] Zhang M, Molinos J G, Zhang X, Xu J. Functional and taxonomic differentiation of macrophyte assemblages across the Yangtze River floodplain under human impacts. Frontiers in Plant Science, 2018, 9: 387. doi: 10.3389/fpls.2018.00387.

